# Galectin-9 has non-apoptotic cytotoxic activity towards Acute Myeloid Leukemia independent of cytarabine resistance

**DOI:** 10.1101/2023.01.12.523722

**Authors:** Ghizlane Choukrani, Nienke Visser, Natasha Ustyanovska Avtenyuk, Mirjam Olthuis, Glenn Marsman, Emanuele Ammatuna, Harm Jan Lourens, Toshiro Niki, Gerwin Huls, Edwin Bremer, Valerie R. Wiersma

## Abstract

Acute myeloid leukemia (AML) is a malignancy still associated with poor survival rates, among others due to frequent occurrence of therapy-resistant relapse after standard-of-care treatment with cytarabine (AraC). AraC triggers apoptotic cell death, a type of cell death to which AML cells often become resistant. Therefore, therapeutic options that trigger an alternate type of cell death are of particular interest. We previously identified that the glycan-binding protein Galectin-9 (Gal-9) has tumor-selective and non-apoptotic cytotoxicity towards various types of cancer, which depended on autophagy inhibition. Thus, Gal-9 could be of therapeutic interest for (AraC-resistant) AML. In the current study, treatment with Gal-9 was cytotoxic for AML cells, including for CD34^+^ patient-derived AML stem cells, but not for healthy cord blood-derived CD34^+^ stem cells. This Gal-9-mediated cytotoxicity did not rely on apoptosis but negatively associated with autophagic flux. Importantly, both AraC-sensitive and -resistant AML cell lines as well as AML patient samples were sensitive to single agent treatment with Gal-9. Additionally, Gal-9 potentiated the cytotoxic effect of DNA demethylase inhibitor Azacytidine (Aza), a drug that is clinically used for patients that are not eligible for intensive AraC treatment. Thus, Gal-9 is a potential therapeutic agent for the treatment of AML, including AraC resistant AML, by inducing caspase-independent cell death.

## Introduction

Acute myeloid leukemia (AML) is an aggressive hematologic malignancy characterized by accumulation of various cytogenetic and molecular alterations [1, 2]. The standard treatment regimen of AML patients comprises the nucleoside analogue cytarabine (AraC) combined with an anthracycline such as daunorubicin or idarubicin. However, prolonged disease-free survival following therapy with AraC is uncommon due to the development of therapy resistance [3]. Consequently, the 5-year overall survival rate for AML is just 25%, although there is a significant variability depending on AML subtype, ranging from 5-10% overall survival for individuals with poor risk AML to over 90% overall survival for patients with acute promyelocytic leukemia [4].

One of the primary mechanisms underlying AraC resistance in AML is evasion of the apoptotic cell death program, with e.g., deregulated expression of proteins that regulate apoptosis among which Bcl-2 and Bcl-xl [5-7]. Thus, novel therapeutic strategies that do not rely on apoptosis are of particular interest for the treatment of relapsed patients with AraC resistant AML. Furthermore, patients that are not eligible for intensive chemotherapy, particularly elderly patients (>65 years of age) and those with intermediate-risk cytogenetics, are treated with the hypomethylating agent azacitidine (Aza) [8]. However, the survival rate of those patients after Aza treatment is merely 14 months [8], making the search for more effective treatment options for these patients warranted.

An interesting candidate in this respect is Galectin-9 (Gal-9), a carbohydrate-binding protein that is comprised of two carbohydrate recognition domains connected by an inter-domain linker [9]. We and others have previously demonstrated that a recombinant form of Gal-9 has potent cytotoxic activity towards various cancer types [10-14]. Furthermore, Gal-9 killed imatinib-resistant chronic myelogenous leukemia cells [18]. Although associated with features of apoptosis, like phosphatidylserine exposure, Gal-9-induced cytotoxicity could not be blocked by pan-caspase inhibition and thus did not require apoptotic signaling [13, 14]. Instead, cytotoxicity associated with the induction and halted execution of its proper execution [14]. Of note, whereas Gal-9 had potent cytotoxic activity toward cancer cells, it did not negatively affect the viability of healthy counterparts [13, 14]. Based on this non-apoptotic cytotoxic activity of Gal-9, we hypothesized that Gal-9 may also be of potential interest for the treatment of AML in general, as well as for therapy resistant AML.

In the current study, treatment with recombinant Gal-9 induced cell death in both AML cell lines and primary patient-derived AML cells, including CD34^+^ AML stem cells, but not healthy CD34^+^ cord blood-derived stem cells. Furthermore, AraC-sensitive as well as AraC-resistant AMLs were equally susceptible to Gal-9 cytotoxicity. Gal-9-mediated cytotoxicity towards AML cells did not rely on apoptotic signaling but was associated with halted execution of autophagy. Finally, the combination of Gal-9 with Aza induced more cell death compared to either Aza or Gal-9 treatment alone. Thus, Gal-9 is a potential novel therapeutic agent for the treatment of AML, including AraC resistant AML.

## Materials and methods

### Cell Lines and Galectin-9

THP-1, HL-60, U-937, MOLM-13, NB4, OCI AML2, OCI AML3 and MS5 were originally obtained from American Type Culture Collection (ATCC). All cell lines were cultured at 37°C, in a humidified 5% CO_2_ atmosphere in RPMI (Lonza 12-115F, Basel, Switzerland) supplemented with 10% fetal calf serum (FCS) (Sigma Aldrich, F7524, St. Louis, MO, USA). Cytarabine-resistant cell lines were generated as described previously by us [15]. Recombinant Galectin-9 (Gal-9) was produced as previously described [16]. AraC and Aza were from the UMCG hospital pharmacy.

### Ex Vivo Culturing of Patient-Derived AML and cord-blood cells

Patient-derived AML samples (stored after informed consent and following approval by the Medical Ethical committee of the UMCG in accordance with the Declaration of Helsinki protocol code NL43844.042.13, 6 January 2014) were thawed from cryovials and added to pre-warmed newborn calf serum (NCS, Gibco, Breda, The Netherlands), and centrifuged at 450g for 5 min. The cell pellet was then resuspended in pre-warmed NCS mix (5 U/mL Heparin (Pharmacy of the UMCG), 4μM magnesium sulphate (Sigma-Aldrich, St. Louis, MO, USA), and 20U/mL DNase (Roche, Basel, Switzerland)) and incubated for 15 min in a 37°C water bath. Thereafter, AML cells were washed and cultured in Gartner’s medium (Alpha-MEM, 12.5% horse serum, 12.5% FCS, 1μM hydrocortisone (Sigma-Aldrich, St. Louis, MO, USA), 1% pen-strep (Sigma-Aldrich, St. Louis, MO, USA), and 50μM beta-mercaptoethanol (Sigma-Aldrich, St. Louis, MO, USA)) supplemented with thrombopoietin (TPO) and G-CSF, IL-3 (20 ng/mL of each cytokine) (hospital pharmacy, UMCG), as described before [17] on top of a MS5 support-layer grown on gelatin coated flasks for 2–3 days. After this recovery period, primary samples with a viability over 80% were used in cell death and autophagy assays. Primary human mesenchymal stem cells (MSCs) were cultured on gelatin coated flasks for 2-3 days in Alpha-MEM (Sigma-Aldrich, St. Louis, MO, USA) supplemented with 1% pen-strep (Sigma-Aldrich, St. Louis, MO, USA), 10 U/mL Heparin (Pharmacy of the UMCG) and 5% platelet lysate (Sigma-Aldricht, St. Louis, MO, USA).

Mononuclear cells (MNCs) were isolated from Cord Blood (CB) of healthy subjects and separated using ficoll (Lymphoprep™, Bernburg, Germany), after which CD34^+^ stem cells were isolated by MACS sorting using CD34 MACS microbeads (Miltenyi Biotec, Leiden, The Netherlands) following manufacturers recommendations, and the negative fraction was used as. CD34^-^ cells. CD34^+^ from patient derived AML samples were isolated in the same way.

### Cytotoxicity Assays

To determine the cytotoxicity of AraC, Aza, Gal-9 and CQ (from the LC3B Antibody Kit for Autophagy, L10382, Invitrogen™, Carlsbad, CA, USA), 5×10^4^ cells (either cell lines or primary patient-derived material) were plated in a 48 well plate with 200μL RPMI supplemented with 10% FCS (for cell lines) or Gartner’s medium (for patient-derived AML cells or CB cells) and treated with the indicated concentrations of Gal-9, AraC, Aza or CQ. In case of combinational treatments, Gal-9 and AraC were added simultaneously, whereas cells were pre-incubated with Aza for 16h before adding Gal-9. For experiments with Z-VAD-fmk (R&D Systems, Inc., FMK001, Wiesbaden, Germany) cells were pre-treated with 20μM Z-VAD-fmk for 16h, and again treated with 20μMZ-VAD-fmk freshly added 1h before adding Gal-9 or Staurosporin (Sigma Aldrich, St. Louis, MO, USA). For long-term assays using patient-derived AML cells (5-7 days), wells were first coated with MS5 cells (pre-plated 1 day before the assay to reach confluency at day of use) before AML cells were layered on top of the stromal layer. For the CQ assays, cells were incubated on top of MS5 cells. Cytotoxicity was assessed using either flow cytometry-based cell counts, Annexin-V staining, DioC6 staining or MTS assay.

### Flow cytometry-based cell count

After 24h of incubation, cells were harvested and counted using a flow cytometer (BD Accuri™ C6 cytometer, BD Biosciences, San Jose, CA, USA or Cytoflex, Beckman Coulter, Brea, CA, USA and accessory software). Here, cells/ μl in the ‘viable’ gate was determined (Figure 1B).

**Figure 1:**
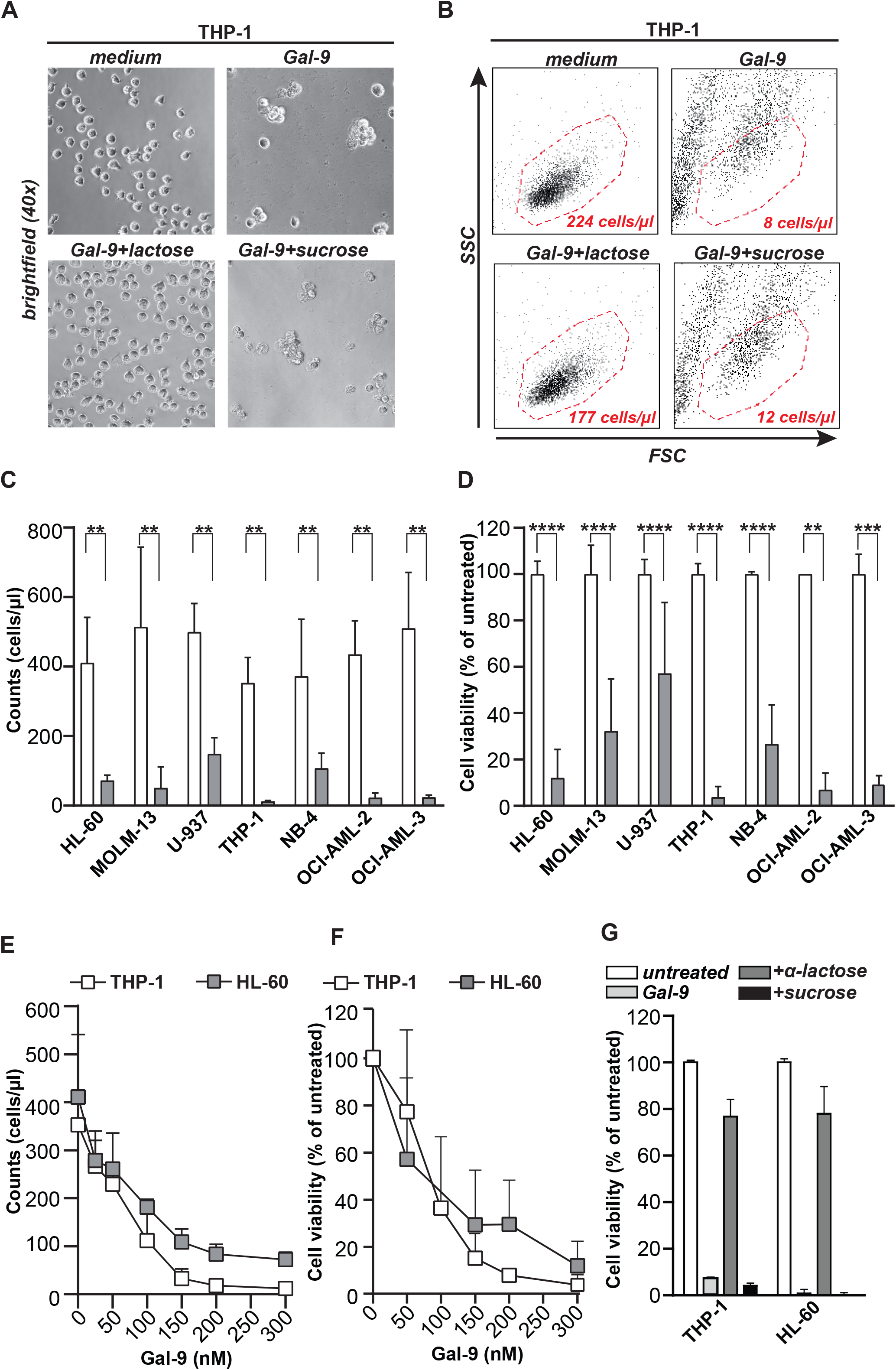
Gal-9 is cytotoxic for AML cell lines. Viability of THP-1 cell line treated with 300nM Gal-9 for 16h, visualized by microscopy **(A)** or **(B)** flow cytometry. **(C)** Flow cytometry-based cell counts or **(D)** cell viability using the MTS assay upon treatment of different AML cell lines with Gal-9 (300nM) for 16 or 72h, respectively. **(E)** Dose dependent impact of Gal-9 on flow cytometry-based cell counts or **(F)** cell viability upon treatment with the indicated concentrations of Gal-9 for 72h. **(G)** Percentage of cell viability upon treatment with 300nM Gal-9 in the presence of α-lactose or sucrose (40mM).

### MTS assay

After 72h of incubation, cell viability was assessed using the MTS assay (CellTiter 96^®^ AQueous One Solution Cell Proliferation, G3580, Promega, Madison, WI, USA). In brief, MTS was added (7.5 % v/v) to each well and incubated at 37°C. After sufficient color development read-out was performed at OD490nm (Multi-Scan Sky of Thermo Scientific Waltham, MA, USA). Cell viability was calculated by subtracting OD490nm of the dead control of each value (7.5 % ‘dead mix’, consisting of 10% Triton-X in 70% ethanol) and calculating the viability as % of the untreated control (treated/untreated*100%).

### Annexin-V assay

After 24h of incubation, cells were harvested and stained with Annexin-V. In brief, cells were resuspended in 1x Annexin V binding buffer (10x Annexin V binding buffer, BD Biosciences, San Jose, CA, USA) and 1% Annexin-V-FITC (Immunotools, Friesoythe, Germany) was added. After 10 minutes incubation at 4°C, staining was analyzed using flow cytometry.

### DioC6 staining

DiOC6 (Molecular Probes, Eugene, Oregon, USA) at a concentration of 0.1 μM in fresh culture medium was added 1:1 to cells. After 20 minutes of incubation at 37°C, cells were collected by centrifugation (450g, 5min), resuspended in PBS, and analyzed using flow cytometry.

To determine the percentage of CD34^+^ AML cells after treatment with Gal-9, AML cells were pre-incubated with FcR-blocker (100μg/mL) (Miltenyi Biotec, Leiden, The Netherlands) and subsequently stained with anti-CD34 antibody ((Clone: 561, Biolegend, California, USA) for 1h at 4°C. Cells were then collected by centrifugation (450g, 5min), resuspended in PBS, and analyzed using flow cytometry.

### Autophagosomal and lysosomal content: Cyto-ID and lysotracker

To determine the autophagosomal and lysosomal content in cell lines and AML patient-derived samples, cells were cultured at a density of 5 × 10^4^ in a 48 wells plate and treated with Gal-9 (300nM) or with CQ (10μM) for 6h. Subsequently, LysoTracker® Red DND-99 (1 μM, Life Technologies, L-7528, Carlsbad, California, USA) or cyto-ID was added in combination with hoechst (Cyto-ID autophagy detection kit; ENZ-51031–0050, Enzo lifesciences, Inc., New York, USA) following the manufacturer’s recommendation, whereupon cells were incubated for 30min at 37°C. Dye excess was washed away with PBS and staining was visualized using IncuCyte S3 Live-Cell Analysis System (Ann Arbor, MI, USA) or the EVOS Cell Imaging System (EVOS-FL, Thermo Scientific, Waltham, MA, USA). The corrected total cell fluorescence (CTCF) of LysoTracker and Cyto-ID signal was determined using ImageJ software.

### Western blot analysis of autophagic flux and caspase-3 activation

To detect the basal level of autophagic flux in AML cells, 1 × 10^6^ cells were cultured in a 6 well plate and treated with CQ (50μM) for 6h or 24h. To determine the impact of Gal-9 on the autophagy pathway, cells were treated with Gal-9 (300nM) for 6h or 24h. Cell lysates were prepared using lysis buffer (50 mM Tris, 2 mM EDTA, 2 mM EGTA, 150 mM NaCl, 0.1% SDS, 1% NP-40 substitute) containing 1μM Na_3_VO_4_ (Sigma, 450243, St. Louis, MO, USA) and protease inhibitor cocktail (Sigmafast; Sigma Aldrich, S8820, St. Louis, MO, USA). Protein concentration was determined using Bradford protein assay (Pierce™ Coomassie (Bradford) Protein Assay Kit, #23200, Thermo Scientific, Waltham, MA, USA) then 20μg of total protein was loaded to SDS-PAGE gels for electrophoresis using a 15% gel. Next, proteins were transferred to a nitrocellulose membrane (Amersham Hybond ECL; GE Healthcare, RPN303D, Merck, Darmstadt, Germany). After blocking in 5% (w/v) milk powder/TBST, proteins were detected using primary antibodies against LC3B (LC3B Antibody Kit for Autophagy, Invitrogen™, L10382, Carlsbad, CA, USA), SQSTM1/ p62 (SantaCruz, sc-28359, California, USA), β-actin (Abcam, ab49900, Cambridge, UK), total caspase 3 (#9665 cell signaling, Danvers, Massachusetts, USA), cleaved caspase 3 (#9661 cell signaling, Danvers, Massachusetts, USA), and appropriate secondary HRP-conjugated antibodies (Dako, p0217, p0260, Santa Clara, USA). Blots were developed using chemiluminescent substrate (SuperSignal West Dura, Thermo Scientific, Life Technologies, 34075, Waltham, MA, USA) and imaged using the ChemiDoc MP system (Bio-Rad, Hercules, California, USA). Quantification of detected protein levels was performed using the ImageJ tool for gel analysis.

### Confocal microscopy

To detect Gal-9 accumulation in lysosomes, THP-1 cells were seeded in a 6 wells plate (5 × 10^5^ cells/mL) and treated with Gal-9-Alexa-594 overnight. Gal-9-Alexa-594 was prepared using DyLight® 594 (DyLight 594 NHS Ester; Piercenet, Thermo Scientific, 46412, Waltham, MA, USA) following manufacturer’s protocol. Subsequently, cells were transferred to a microscope slide using a cytocentrifuge (Cytospin 3, Shandon, England). 4% w/v paraformaldehyde (PFA) was used for fixation for 15min, and then the lysosomes were stained with anti-LAMP-1-488 (sc-20011 AF488 Santacruz Biotechnology, Dallas, Texas, USA) in culture medium containing 40mM α-lactose to prevent any direct interaction of Gal-9 with the LAMP-1 antibody. After 1h incubation at RT, the cells were washed twice with TBST, and nuclei were stained with DAPI in a mounting medium (Sigma Aldrich, D9542, St. Louis, MO, USA) and subsequently visualized using a Leica SP8 Confocal microscope (Leica Microsystems, Rijswijk, The Netherlands).

### Statistical Analysis

EC50 and sigmoidal curve fitting correlation coefficients of Gal-9 were calculated using the ED50 Plus v1.0 Excel worksheet developed by Dr. Mario H. Vargas at Instituto Nacional de Enfermedades Respiratorias. Significance was tested using Wilcoxon signed-rank test (treated vs. untreated in the same sample) or in case the samples were not ranked (different samples in the groups) using the Mann-Whitney U test using Graphpad Prism software (GraphPad Prism; GraphPad Software, La Jolla, CA, USA). p values are indicated as: **** *p* < 0.0001, *** *p* < 0.001, ** *p* < 0.01, and * *p* < 0.05.

## Results

### Galectin-9 is cytotoxic for AML cell lines

Previously, we identified that treatment of various solid cancers with recombinant Gal-9 induced cancer-specific cell death [13, 14]. In line with this finding, ‘short-term’ treatment of a panel of AML lines with Gal-9 (300nM) strongly and significantly reduced the number of viable cells, as visualized using microscopy for THP-1 (Figure 1A). Upon quantification using flow cytometry, Gal-9 treatment strongly reduced viable THP-1 cell counts compared to control (Figure 1B), as well as an extended AML cell line panel (Figure 1C). This effect was dose-dependent (Suppl. Figure 1A-E) with EC50 for cell lines ranging from 92-193nM for cell count (Suppl. Figure 1K). Correspondingly, treatment with Gal-9 significantly and dose-dependently reduced cell count and viability THP-1 and HL60 cells (Figure 1D-F and Suppl. Figure 1F-J), with EC50s ranging from 115-300nM for viability (Suppl. Figure 1K). This cytotoxic activity of Gal-9 was inhibited by co-incubation with the carbohydrate recognition domain (CRD)-blocking sugar α-lactose, but not by the irrelevant sugar sucrose, demonstrating CRD-dependent activity (Figure 1G and Suppl. Figure 1L, M). Thus, Gal-9 had direct and dose-dependent cytotoxic activity towards AML cells.

#### Galectin-9 is cytotoxic for patient-derived CD34^+^ AML stem cells and CD34^-^ AML blasts, but not for CD34^+^ cord blood-derived stem cells

The cytotoxicity of Gal-9 was further assessed towards primary patient-derived AML cells, with clear reduction in cell density in both unsorted, CD34^+^ AML stem cell and CD34^-^ AML blast populations after treatment with Gal-9 (Figure 2A). This effect was again CRD-specific as α-lactose abrogated cytotoxicity (Suppl. Figure 2A) and was dose-dependent (Figure 2B-D). Notably, cytotoxicity towards CD34^+^ AML cells was slightly higher than towards CD34^-^ AML, albeit not significantly (Suppl Figure 2B). Nevertheless, treatment with a low dose of Gal-9 reduced the percentage of CD34^+^ cells in a mix culture with CD34^-^ cells (Suppl Figure 2C, D).

**Figure 2:**
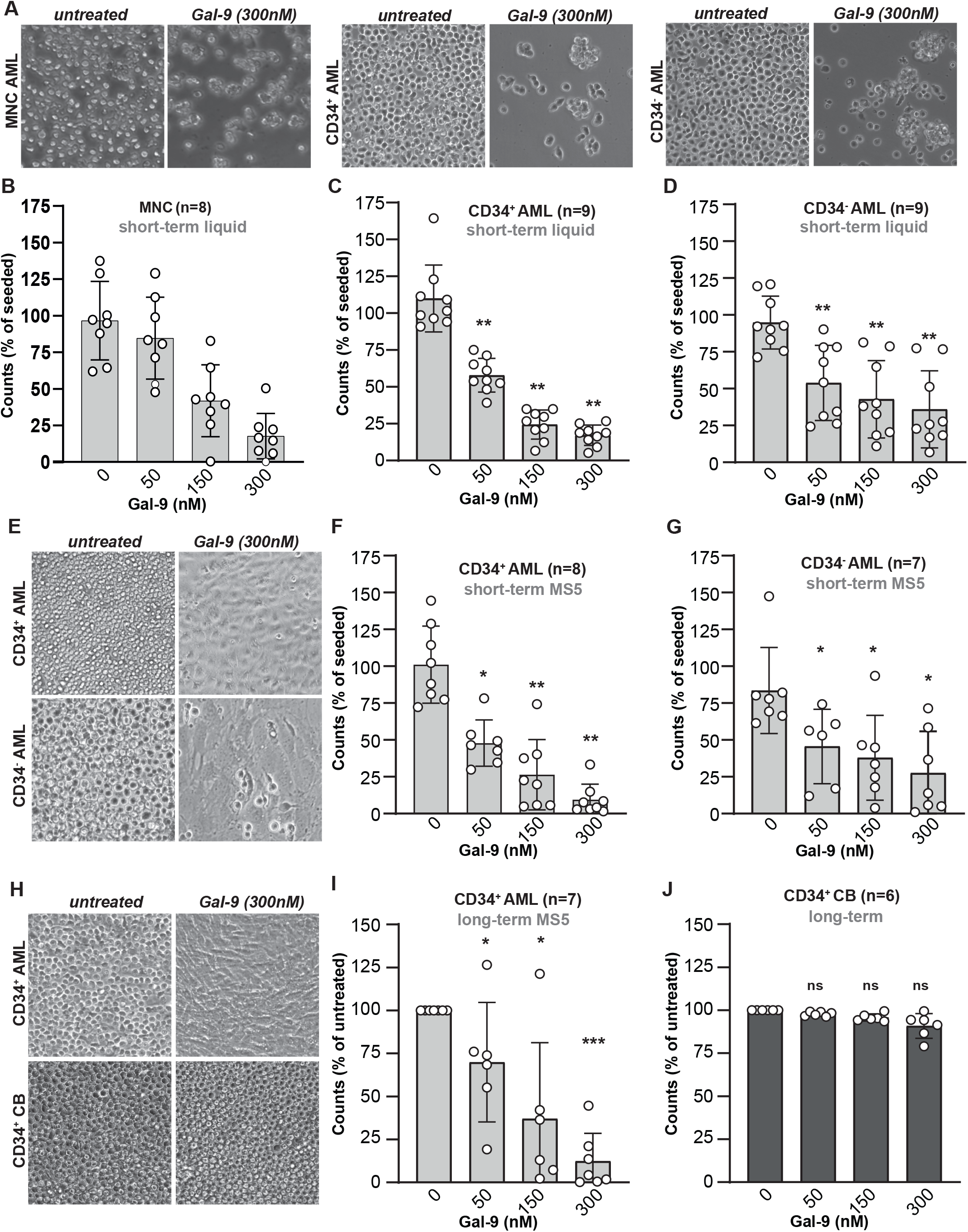
Gal-9 is cytotoxic for patient-derived AML cells, but not CD34^+^ cord blood-derived cells. **(A)** Microscopic images of Gal-9-treated patient-derived AML including MNC AML, CD34^+^ AML and CD34^-^ AML cells. Dose-dependent cytotoxicity of Gal-9 in liquid culture (1-3 days) for patient-derived AML cells in either the **(B)** MNC AML, **(C)** CD34^+^ AML, or **(D)** CD34^-^ AML cell fraction. **(E)** Microscopic picture of patient-derived CD34^+^ and CD34^-^ AML cells treated with 300nM Gal-9 (16h) on top of a MS5 stromal layer. Short-term (16h) treatment of **(F)** patient-derived CD34^+^, or (G) CD34^-^ AML cells on a MS5 stromal layer using the indicated concentrations of Gal-9. **(H)** Microscopic pictures of patient-derived CD34^+^ AML and CB-derived CD34^+^ cells treated with 300nM Gal-9 (6 days) on a MS5 stromal layer. Long-term incubation (5-7 days) of **(I)** CD34^+^ AML or **(J)** CD34^+^ CB cells on top of a MS5 stromal layer with the indicated doses of Gal-9 and evaluated using flow cytometry-based counts.

In the bone marrow niche AML cells are surrounded and supported by stromal cells, with in vitro co-cultures typically being less sensitive to treatment [18]. However, short-term treatment of AML/MS5 stromal cell co-cultures with Gal-9 almost complete eliminated CD34^+^ as well as CD34^-^ AML cells from co-cultures, with only a monolayer of unaffected MS5 remaining (Figure 2E, see also data in figure 4). In line with this, Gal-9 treatment of MS5 cells alone did not impact on viability (Suppl. Figure 2E). In a panel of patient-derived AML cells in co-culture with MS5, Gal-9 dose-dependently reduced viability by ∼90% for CD34^+^ AML and ∼75% for CD34^-^ AML, respectively (Figure 2F, G). No significant difference in sensitivity was detected between the AML stem cell and blast population (Suppl. Figure 2F), although the cytotoxic effect of Gal-9 in AML/MS5 cocultures was significantly higher than in liquid cultures without stromal support (Suppl. Figure 2G). Again, Gal-9 cytotoxicity was abrogated by co-incubation with α-lactose (Suppl. Figure 2A, H).

Importantly, treatment with a single dose of Gal-9 eliminated the CD34^+^ AML cells even after ‘long-term’ incubation of up to 7 days, whereas cord blood-derived CD34^+^ stem cells remained unaffected as shown by microscopy images (Figure 2H). The cytotoxic effect of Gal-9 on CD34^+^ AML cells in these longer-term co-cultures was dose-dependent (Figure 2I), not affecting CD34^+^ cord blood cells at any concentration (Figure 2J), yielding the strongest differential effects at 300nM Gal-9 (Suppl. Figure 2I). In addition, whereas Gal-9 reduced mitochondrial membrane potential as measured by DioC6 in CD34^+^ AML cells, this effect was not observed in CD34^+^ cord blood cells (Suppl. Figure 2J). Like for CD34^+^ AML cells, Gal-9 dose-dependently eliminated CD34^-^ AML cells in long-term assays (Suppl. Figure 2K). Notably, repeated treatment with a low dose of Gal-9 (25-50nM) every 3^rd^ day yielded a similar reduction in cell counts as a single dose with 300nM Gal-9 after 2 weeks (Suppl. Figure 2L). Taken together, Gal-9 had a dose-dependent cytotoxic effect towards patient-derived CD34^+^ as well as CD34^-^ cells in both liquid and MS5 co-cultures, whereas it did not negatively impact on MS5 stromal cells and healthy cord blood-derived CD34^+^ stem cells.

### Galectin-9-induced AML cell death did not rely on caspase-dependent apoptosis

In several reports, Gal-9-induced cell death was reported to rely on apoptotic signaling [19-21]. In contrast, we previously demonstrated that Gal-9-induced cell death, although being characterized by the hallmark apoptosis feature of phosphatidylserine (PS)-exposure, was caspase-independent in colon cancer and melanoma cells [13, 14]. Also in AML, PS-exposure was detected upon Gal-9 treatment of THP-1 cells and CD34^+^ patient-derived AML cells in both liquid and MS5 co-cultures (Figure 3A-C). However, this effect was not blocked by co-incubation with the pan-caspase inhibitor Z-VAD-FMK in cell lines (Figure 3D) nor patient-derived AML cells (Figure 3E). In contrast, Z-VAD-FMK did reduce PS-exposure upon treatment with apoptosis-inducer staurosporine (Figure 3D). Furthermore, no processing of caspase-3 was detected in Gal-9-treated HL-60 cells, whereas staurosporine did induce caspase-3 processing (Figure 3F). Like the reduction in cell counts and cell viability, PS-exposure was dose-dependent (Suppl. Figure 3A-G), with an EC50 ranging from 77-140nM in the cell line panel (Suppl Figure 3H), and CRD-dependent (Figure 3G-I). PS-exposure was less pronounced in MS5 co-cultures and especially in CD34^-^ AML cells, only reaching significance at 300nM Gal-9 (Suppl. Figure 3I-L). Furthermore, Gal-9 treatment did not induce PS-exposure in CD34^+^ cord blood cells (Suppl. Figure 3M). Thus, although Gal-9 is a potent inducer of PS-exposure, Gal-9-induced cell death does not require caspase-activation.

**Figure 3:**
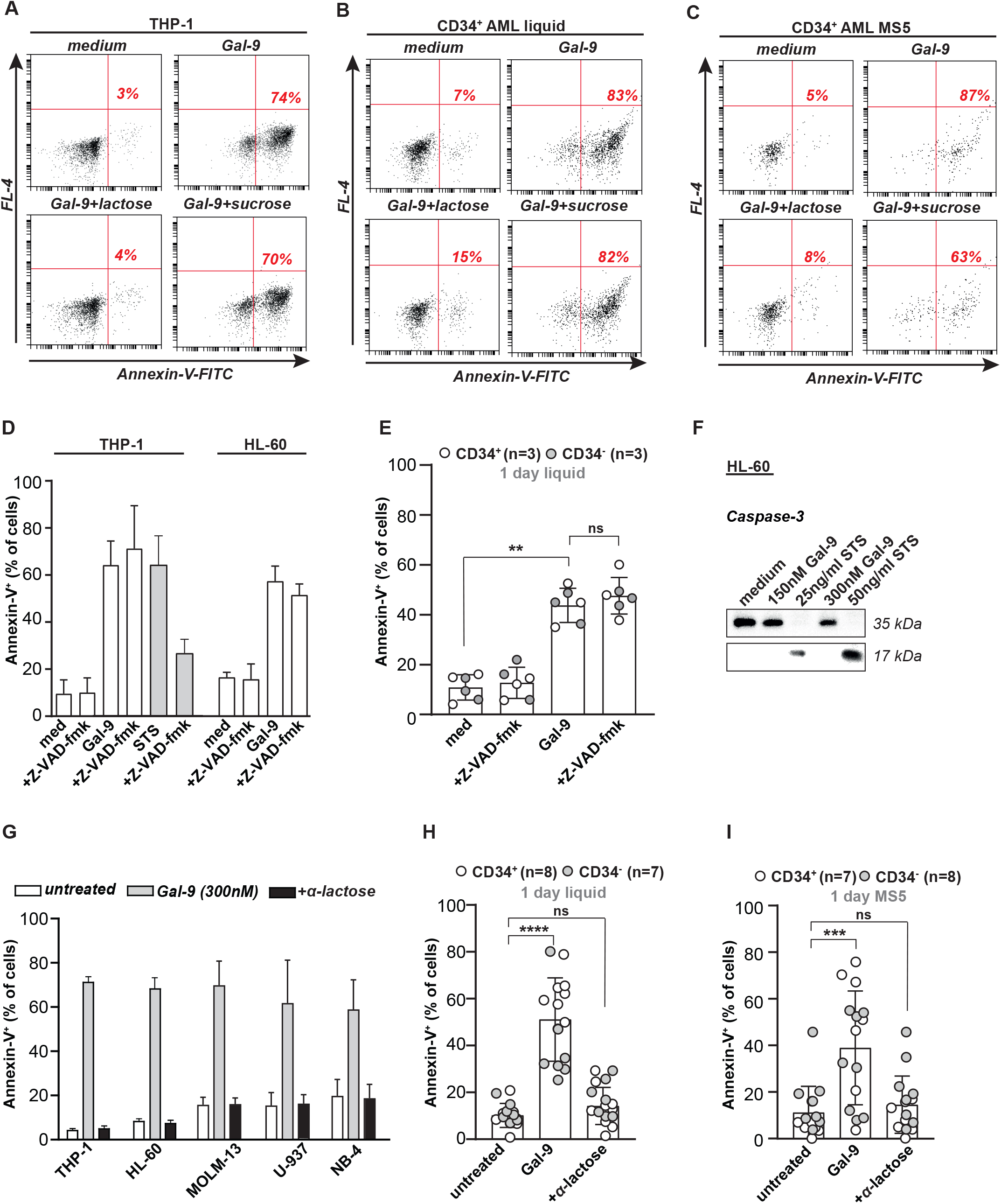
Gal-9-induced AML cell death does not rely on caspase-dependent apoptosis. PS-exposure detected by flow cytometry in Gal-9-treated (300nM, 16h) **(A)** THP-1 cells, CD34^+^ patient-derived AML cells in **(B)** liquid, or **(C)** MS5 co-cultures. **(D)** Quantification of PS-exposure after blockade of pan-caspases with Z-VAD-FMK in Gal-9-treated THP-1 and HL-60 cells, using staurosporine (STS) as a positive control. **(E)** As in **(D)** but determined on patient-derived CD34^+^ and CD34^-^ cells. **(F)** Western blot of full-length caspase-3 (35 kDa) and cleaved caspase-3 (17 kDa) in HL-60 treated with Gal-9 (150, 300nM, 16h) or STS (25, 50ng/ml, 16h). Blockade of Gal-9-mediated PS-exposure with α-lactose (40mM) in **(G)** a panel of AML cell lines, **(H)** CD34^+^/CD34^-^ patient-derived AML cells in liquid culture, or **(I)** in co-culture with MS5.

### Galectin-9 inhibits the execution of autophagy in AML cells

In colon carcinoma we identified that Gal-9-induced cell death was characterized by prominent vacuolization and depended on the autophagy pathway [14]. Analogously, Gal-9 treatment of AML cells triggered prominent vacuolization, a hallmark of autophagy, as illustrated for THP-1 and CD34^+^ AML cells, which was not detected in healthy CD34^+^ cord blood stem cells (Figure 4A). These vacuoles were partly characterized as autophagosomes with a clear increase in Cyto-ID staining in both cell lines (Figure 4B, C) and CD34^+^ patient-derived AML cells (Figure 4D). In addition, accumulation of lysosomes was detected, whereby Gal-9 triggered a strong increase in Lysotracker staining in AML cell lines (Figure 4E-F) and CD34^+^ patient-derived AML samples (Figure 4G).

**Figure 4:**
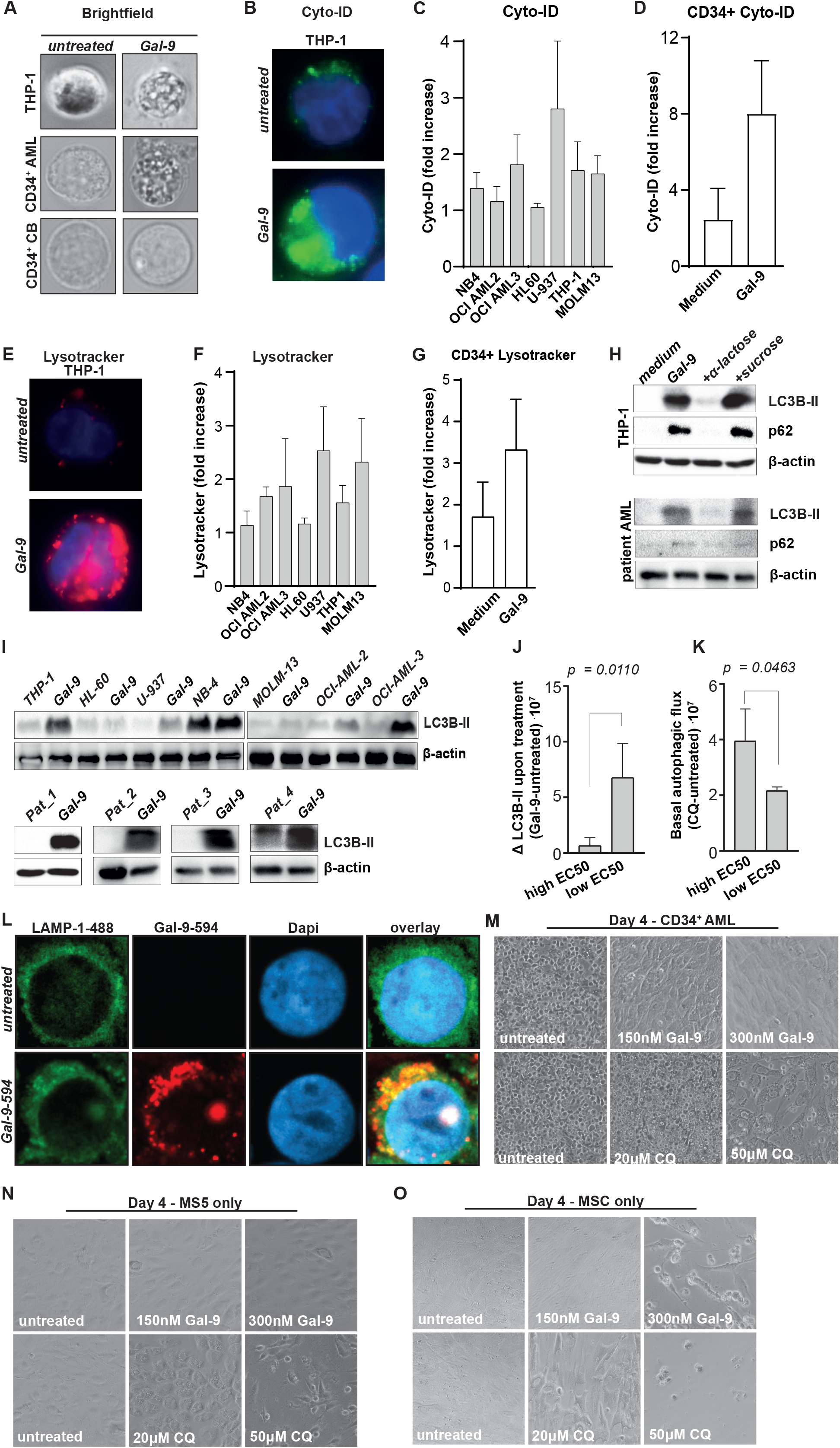
Gal-9 inhibits autophagy in AML cells. **(A)** Microscopic images of AML cells (THP-1 and patient-derived CD34^+^ cells) and CB-derived CD34^+^ cells upon treatment with Gal-9 (300nM, 16h). (B) Fluorescent microscopic image of THP-1 cells upon treatment with Gal-9 (300nM, 16h) and stained with Cyto-ID. **(C)** As in (B) but quantified the fluorescent signal using ImageJ for the AML cell line panel. **(D)** as in (C) but using CD34^+^ patient-derived AML cells. **(E)** Fluorescent microscopic image of THP-1 cells upon treatment with Gal-9 (300nM, 16h) and stained with lysotracker. (F) As in (E) but quantified the fluorescent signal using ImageJ for the AML cell line panel. (G) as in (F) but using CD34^+^ patient-derived AML cells. **(H)** Western blot of autophagy related proteins SQSTM1/p62 (62 kDa) and LC3B-II (16 kDa), including loading control beta-actin (42 kDa), upon incubation with Gal-9 (300nM, 6h) in the THP-1 cell line and a patient-derived CD34^+^ AML sample. **(I)** Western blot of the whole cell line panel as well as 4 different patient-derived AML samples detecting LC3B-II (16 kDa) and the loading control beta-actin (42 kDa) upon treatment with Gal-9 (300nM, 16h). **(J)** Association of the Gal-9 sensitivity (either low EC50/ highly sensitive or high EC50/ weakly sensitive) with the accumulation of LC3B-II upon treatment with Gal-9 as shown in Figure 4I. **(K)** As in (J), but comparing sensitivity for Gal-9 with the basal autophagic flux as determined by incubation with CQ (Suppl. Figure 4B). **(L)** Representative confocal images of Gal-9 accumulation in the lysosomes of AML cells using fluorescently labeled Gal-9 (Gal-9-594; red) and counter stained for LAMP-1 (LAMP1-488; green), and dapi (blue). **(M)** Microscopic pictures of CD34^+^ patient-derived AML cells treated with Gal-9 (150, 300nM) and CQ (20, 50μM), for 4 days on top of a MS5 monolayer. **(N)** Microscopic image of MS5 cells only, treated with the indicated concentrations of Gal-9 and CQ for 4 days. **(O)** As in **(N)** but using primary human mesenchymal stem cells (MSC).

To further investigate the involvement of the autophagy pathway, the hallmark markers LC3B-II and p62/SQSTM1 were assessed. LC3B-II is produced during the activation of autophagy to form the autophagosome, whereas p62 is a cargo protein that helps to entrap target material in the autophagosomes [22]. During the execution phase of autophagy, when the autophagosome fuses with the lysosome, p62 and most of the LC3B-II is degraded. Therefore, persistent accumulation of LC3B-II and p62 is a sign of halted autophagy. Gal-9 induced a CRD-dependent increase in both LC3B-II and p62 levels in AML cell lines (Figure 4H, Suppl. Figure 4A) as well as in a patient-derived CD34^+^ AML sample (Figure 4H) upon 6h of incubation. This accumulation in LC3B-II was persistent in both AML cell lines and patient-derived AML cells as it retained after 24h of incubation (Figure 4I). Notably, the extend of LC3B-II accumulation was not equal in all AML cell lines and associated with the sensitivity of the cell line for Gal-9. Specifically, the accumulation in LC3B-II was significantly stronger in cell lines with a higher sensitivity for Gal-9 (and thus a lower EC50) compared to cell lines that were less sensitive (Figure 4J and Suppl. Figure 4B-C). Furthermore, the basal level of autophagic flux, as identified by accumulation of LC3B-II upon treatment with the lysomotrophic agent chloroquine (CQ) (Suppl. Figure 4B), significantly associated with sensitivity for Gal-9. Here, cell lines with a low basal autophagic flux were more sensitive for Gal-9 (Figure 4K and Suppl. Figure 4D).

The above-described data indicate that Gal-9 disturbs the proper execution of autophagy at the stage of lysosomal-autophagosomal fusion, possibly by accumulating in and impairing the functioning of either of these organelles. In line with this, Gal-9 clearly accumulated in the lysosomes of AML cells (Figure 4L), suggesting that Gal-9 acts as a lysomotrophic agent. Interestingly, recent studies highlight that targeting the lysosomes may be a potential novel therapeutic strategy for the treatment of AML [24-27], with CQ being the agent of choice in most studies. Therefore, the cytotoxic potential of CQ and Gal-9 was compared, with 150nM and 300nM Gal-9 (Suppl. Figure 4E) and 20μM and 50μM CQ (Suppl. Figure 4F) being defined as optimal concentrations to respectively eliminate 50% and 90% of cells on average in cell line co-cultures with MS5. In patient-derived AML cells treatment with 150nM Gal-9 was sufficient to achieve 90% cell death after 16h, and complete removal after 4 days of treatment. In contrast, 50μM CQ reduced the viability by only ∼40%, reaching up to 80% after 4 days of treatment (Figure 4M, Suppl. Figure 4G).

Importantly, whereas the MS5 stromal layer underneath Gal-9-treated AML cells remained healthy, this layer was strongly affected upon CQ-treatment (Figure 4M). In line with this, Gal-9 did not negatively impact on the cell viability of MS5 in single culture, whereas the effective dose of CQ for AML cell killing (50μM) had a clear negative impact on MS5 cells (Figure 4N). To further investigate the safety profile of Gal-9 and CQ for healthy cells, primary human mesenchymal stem cells (MSCs) were also treated. Gal-9 did not affect MSC viability at 150nM, whereas CQ at 20 μM already significantly reduced viability (Figure 4O). Furthermore, the MSC monolayer was completely eradicated when treated with 50 μM CQ, whereas, although clearly affected, a substantial amount of MSCs remained adhered to the plate upon treatment with 300nM Gal-9 (Figure 4O).

Taken together, Gal-9 inhibited the proper execution of autophagy by accumulating in the lysosomes and seems to act as a lysomotrophic agent like CQ. However, although both agents are capable of inducing AML cell death, Gal-9 seemed to be less toxic for stromal cells than CQ and therefore may be a more suitable candidate for the treatment of AML.

### Gal-9 is cytotoxic for AraC-resistant AML and potentiates the efficacy of Azacitidine

AraC-resistant (AraC-Res) AMLs often have defects in the apoptosis pathway [23]. Furthermore, lysosomes are known to contribute to therapy resistance by drug sequestration [24]. Since Gal-9 induces apoptosis-independent cell death and acts as a lysosomal inhibitor, we hypothesized that Gal-9 may retain cytotoxicity against AraC-Res AML cells. Indeed, Gal-9 eliminated all four AraC-Res AML cell lines in a dose-dependent manner (Figure 5A-D). The EC50 in HL-60 cells did not differ between parental and AraC-Res cells, slightly increased in AraC-Res MOLM-13 and THP-1 cells, but strongly reduced in U-937 cells (Figure 5A-D). Gal-9 also increased the amount of cell death when combined with AraC treatment compared to either agent alone in parental U-937 cells (Suppl. Figure 4H). Furthermore, patient-derived AML cells that did not respond to AraC were eliminated by Gal-9 in a dose-dependent manner, to a similar extent as samples that were sensitive to AraC (Suppl. Figure 4I, J and Figure 5E, F). Interestingly, in a unique matched *de novo* and relapsed patient sample pair, Gal-9 efficiently killed patient-derived AML cells during both initial disease and during relapse (Figure 5G, 5H).

**Figure 5:**
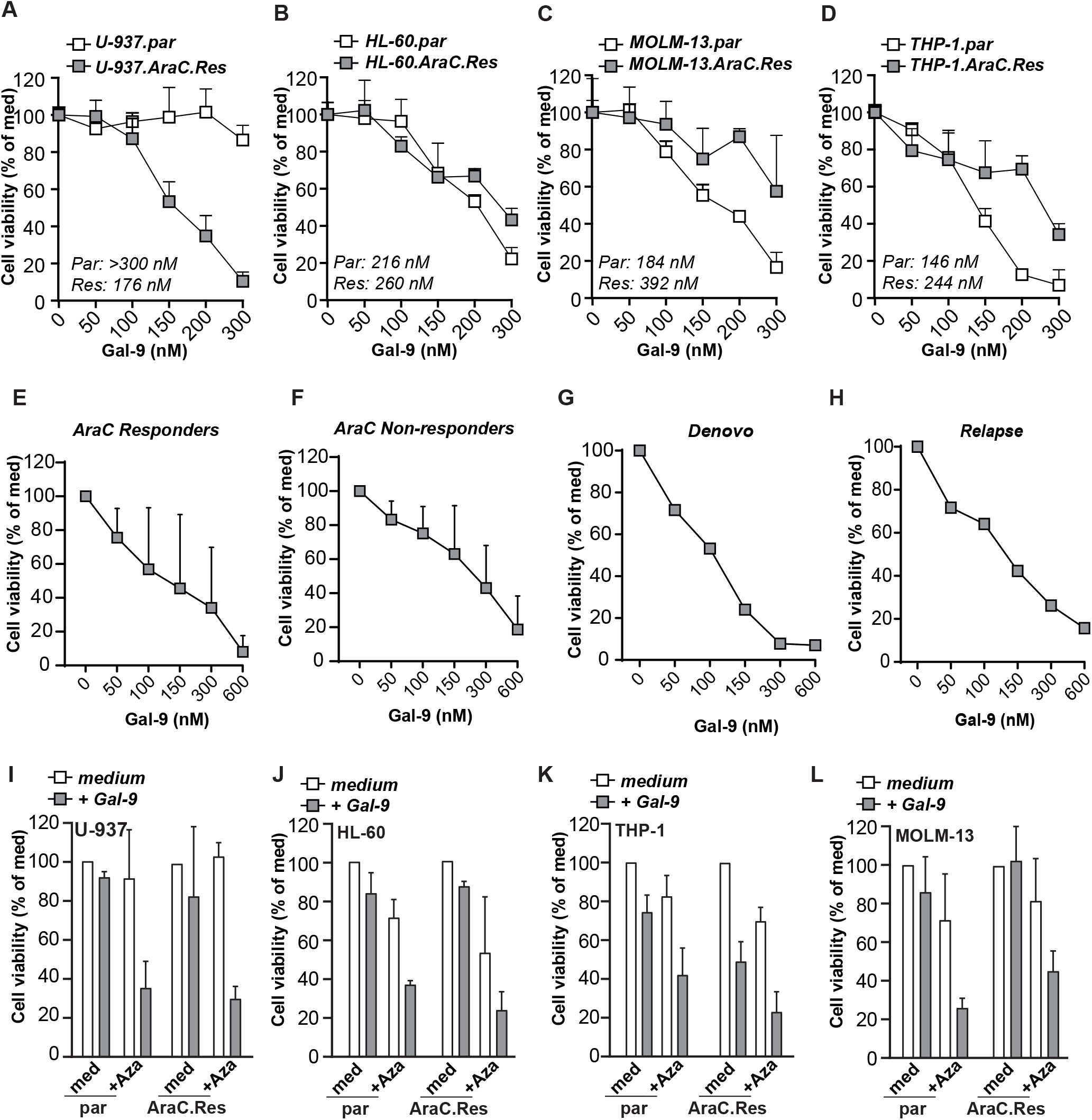
Gal-9 is cytotoxic for AraC-resistant AML and potentiates the efficacy of Aza. Cell viability as determined by the MTS assay of parental vs. AraC resistant (AraC-Res) AML cell line pairs **(A)** U-937, **(B)** HL-60, **(C)** MOLM-13 and **(D)** THP-1, treated with the indicated concentration Gal-9 for 72h. Concentrations depicted inside the graph indicate the EC50 for Gal-9. Impact of Gal-9 as determined by flow cytometry-based cell viability (Annexin V/PI) on **(E)** AraC-responders vs. **(F)** AraC-non responders after 16h of incubation. **(G)** as in (E) and (F), but for a matched sample during initial disease (de novo; **G**) or relapse **(H)**. Cell viability (using MTS assay) of the parental vs. AraC resistant AML cell line panel, including (I) U-937, **(J)** HL-60, **(K)** THP-1, and (L) MOLM-13, treated with a combination of low dose Aza (2,5μM) and Gal-9 (150nM). Of note, Aza was pre-incubated for 16h before adding Gal-9 and incubating for an additional 72h.

Interestingly, combination of Aza treatment with low dose of Gal-9 potentiated the induction of cell death in parental AML cell lines (Figure 5I-L). Furthermore, Gal-9 also potentiated the efficacy of Aza in AraC-resistant AML cells (Figure 5I-L), whereas the combination AraC and Aza was only effective in parental cell lines (Suppl. Figure 4K-N). Thus, the addition of Gal-9 to Aza therapy may be of interest to achieve higher response rates in AML patients that are ineligible for AraC treatment, or patients that relapsed after initial AraC therapy.

## Discussion

In the current study, we identified that Gal-9 induced caspase-independent cell death in AML cell lines and patient-derived AML cells yet did not affect healthy cord blood-derived CD34^+^ stem cells. This induction of cell death was associated with accumulation of Gal-9 in lysosomes and inhibition of the execution phase of autophagy. Importantly, Gal-9 cytotoxicity was retained in AraC-resistant AML cells and potentiated the efficacy of Aza. This sensitivity of AraC-resistant AMLs to Gal-9 cytotoxicity is of potential clinical relevance since new treatment modalities that could surmount apoptosis resistance in AML cells are urgently needed.

As shown in this study, treatment with Gal-9 resulted in halted autophagy and concomitant cytotoxic elimination of AML cells. This finding is similar to our previous study in colorectal cancer [14], which implies that Gal-9 has a unique cytotoxic potential in different cancer types. Autophagy is an essential catabolic process which has been associated with survival and homeostasis in cancer. For instance, increased levels of autophagy were reported in resistant AML cells upon treatment with AraC [15, 25-28], and other drugs like hDAC and mTOR inhibitors [29, 30]. Therefore, agents that inhibit autophagy have been explored for the treatment of AML, which uniformly demonstrated that autophagy inhibition during initial treatment increases treatment efficacy [15, 25-28]. However, timing is an important issue since we recently demonstrated that AML cells that already gained resistance to AraC could not be re-sensitized with autophagy inhibitors [15]. Interestingly, in the current study both AraC sensitive and AraC resistant AML cells were Gal-9 sensitive. This suggest that besides autophagy inhibition, Gal-9 may have additional working mechanisms, not yet elucidated in AML. Potential mechanisms of action are via the induction of reactive oxygen species or via the calcium calpain pathway, as has been respectively demonstrated in ovarium carcinoma [21] and T cell leukemia [31]. Interestingly, endogenous Gal-9 has been reported to regulate autophagy by interacting with lysosome-associated membrane protein 2 (LAMP-2) and, hence, was found to be enriched in the lysosomes [32]. Correspondingly, we observed that exogenously added Gal-9 accumulated in the lysosomes of AML cells, which was in line with our previous study [14]. Whether this accumulation is also the result of Gal-9 interacting with LAMP-2 is still subject of investigation, but the accumulation of Gal-9 in lysosomes suggests that Gal-9 act as lysomotrophic agent.

In AML and solid malignancies, lysosomes are known to sequester chemotherapeutic agents, thereby impacting on their therapeutic efficiency and governing resistance [33]. Therefore, the use of lysomotrophic agents has been proposed to increase treatment efficacy during chemotherapy, with CQ being the most commonly used agent [34-37]. However, the clinical use of CQ or its derivative hydroxychloroquine is hampered by dose limiting toxicities, inducing among others hypoglycaemia and bone marrow suppression, at doses not impacting on the cancer cells [37]. In line with this data, CQ was cytotoxic for both the stromal cell line MS5 and human MSCs at a dose that only affected ∼50% of the patient-derived AML cells in the current study. In contrast, Gal-9 did not negatively impact on MS5 cells, even at the highest dose, and had a less detrimental effect on human MSCs, whilst killing the majority of the patient-derived AML cells. Furthermore, Gal-9 also seems to have a better therapeutic window in stem cells than CQ, as CB-derived healthy CD34^+^ stem cells were equally sensitive to treatment with CQ as AML stem cells [38], whereas Gal-9 did not have any negative impact on CB-derived healthy CD34^+^ stem cells in the current study. Of note, Gal-9 also reduced the percentage of CD34^+^ patient-derived AML cells, although there was no significant difference in sensitivity between CD34^+^ and CD34^-^ cells. This suggest that Gal-9 may induce differentiation of AML stem cells, which is in line with its potential to drive differentiation of various other cell types among which immune cells [39-41] and osteoblasts [42]. Furthermore, Gal-9 also did not have any toxic effects towards healthy melanocytes or colon epithelial cells in our previous studies [13, 14]. Therefore, Gal-9 may be a better option than CQ to block lysosomal functioning, having less cytotoxic side effects towards healthy stromal cells, CD34^+^ stem cells and epithelial cells.

Despite the potent anticancer activity of Gal-9 and seemingly low impact on healthy cells, it is important to take the pleiotropic activity of this lectin towards immune cells into account. On the one hand, Gal-9 is capable of activating immune cells and, thereby, eliciting an anti-cancer immune response. Indeed, we demonstrated that Gal-9 activates neutrophils, resulting in increased cytokine secretion, migration, and survival *in vitro* [43]. Furthermore, we and others delineated that a low dose of Gal-9 activates T cells and triggers their expansion [31, 44], which, although not formally proven yet, may drive anti-cancer immune responses. On the other hand, Gal-9 is known to induce apoptosis at higher concentrations in T-helper 1 and T-helper 17 cells [45], and its expression is associated with T cell effector dysfunction in the tumor microenvironment [46]. Furthermore, Gal-9 has been reported to drive self-renewal of AML stem cells and leukemic progression [46], although this effect was observed at a concentration of 500 pg/ml Gal-9, which is 2.10^4^ times lower than the 300nM we used in our study and that eliminated the fast-majority of CD34^+^ AML stem cells. However, the negative impact of Gal-9 on T cells is a potential limitation to treat patients with Gal-9. One possibility to safely incorporate Gal-9 treatment into clinical practice would be during hematopoietic stem cell transplantation [47]. During this therapy, AML patients receive a high dose of chemotherapy to eradicate all malignant cells. At this stage, Gal-9 can be administered at a high dose, whereby the negative impact on T cells would be of no consideration, as they will be replenished after transplantation. Furthermore, as we demonstrated, Gal-9 increases the efficacy of Aza and also AraC in the U-937 cell line, which suggests that Gal-9 can be added to salvage therapy prior to stem cell transplantation.

In conclusion, Gal-9 has potent cytotoxic activity towards AML cells, but not towards healthy cord blood-derived cells and stromal cells, that is not hampered by AraC resistance. Therefore, Gal-9 is a potential novel therapeutic agent for AML patients in general as well as patients with AraC resistant relapses.

## Figure legends

**Suppl. Figure 1: Gal-9 is cytotoxic for AML cell lines**.

**(A-E)** Flow cytometry-based cell counts for the different AML cell lines upon treatment with the indicated concentrations of Gal-9 (16h incubation). **(F-J)** Cell viability as determined by the MTS assay for the different AML cell lines upon treatment with the indicated concentrations of Gal-9 (16h incubation). **(K)** As in **(A-E)**, but in the presence of α-lactose (40mM) or sucrose (40mM) using 300nM Gal-9. **(L)** As in **(F-J)**, but in the presence of α-lactose (40mM) or sucrose (40mM) using 300nM Gal-9. **(M)** EC50 for Gal-9 as determined based on the concentration curves as depicted in **(A-J)**.

**Suppl. Figure 2: Gal-9 is cytotoxic for patient-derived AML cells**.

**(A)** Microscopic pictures of CD34^+^ patient-derived AML cells treated with Gal-9 (300nM) in liquid culture or on top of a MS5 support layer in the presence or absence of the CRD-blocking sugar α-lactose (40mM). **(B)** Analysis to determine the difference in sensitivity of CD34^+^ vs. CD34^-^ patient-derived AML cells for Gal-9 (300nM) treatment in short-term (16h) liquid culture (cell count based). **(C)** Flow cytometry histogram of untreated (medium) vs. Gal-9-treated (50nM) patient-derived AML cells stained for the stem cell marker CD34 after 3 days of incubation, and **(D)** quantification hereof. **(E)** Representative bright field microscopy pictures of Gal-9 treated MS5 cells. Flow cytometry-based cell counts upon short-term treatment (16h) with Gal-9 (300nM) of (F) CD34^+^ vs. CD34^-^ patient-derived AML cells in MS5 co-cultures, **(G)** CD34^+^ and CD34^-^ patient-derived AML cells in liquid vs. MS5 co-culture, and **(H)** CD34^+^ patient-derived AML samples in short term liquid or MS5 co-cultures in the presence of the CRD-blocking sugar α-lactose (40mM). **(I)** Flow cytometry-based cell counts of long-term incubated healthy cord-blood (CB)-derived CD34^+^ cells vs. patient-derived CD34^+^ AML cells treated with different concentrations of Gal-9. **(J)** Flow cytometry histogram of CB-derived CD34^+^ cells vs. patient-derived CD34^+^ AML cells stained with DioC6 upon treatment with Gal-9 (300nM, 72h). **(K)** Flow cytometry-based cell counts of patient-derived CD34^-^ AML cells treated with a dose range of Gal-9 for long term culture on top of MS5 support cells (5-7 days**). (L)** Flow cytometry-based cell counts of a patient-derived CD34^+^ AML sample after the indicated days of incubation, whereby the cells were treated with Gal-9 every 3 days. Cell count of CD34^+^ patient-derived AML cells after repeated treatment with Gal-9 every 3 days for 14 days.

**Suppl. Figure 3: PS-exposure upon treatment with Gal-9 in AML**

**(A-G)** Detection of PS-exposure on a panel of AML cell lines using flow cytometry-based Annexin-V staining after 16h of incubation using the indicated concentrations Gal-9. **(H)** EC50 for Gal-9 as calculated using the data in (A-G). Detection of PS-exposure using flow cytometry-based Annexin-V staining on patient-derived **(I)** CD34^+^ or (J) CD34^-^ AML cells upon incubation with Gal-9 for 16h in liquid cultures. **(K, L)** As in (I, L) but upon incubation with Gal-9 for 16h in MS5 co-cultures. **(M)** Detection of PS-exposure using flow cytometry-based Annexin-V staining on patient-derived CD34^+^ AML vs. CB-derived CD34^+^ healthy stem cells.

**Suppl. Figure 4: Gal-9 inhibits autophagy in AML cells**.

**(A)** Western blot detection of LC3B-II (16 kDa) in AML cell lines treated with Gal-9 (300nM, 6h) in the presence or absence of α-lactose and sucrose (40mM), using beta-actin (42 kDa) as loading control. **(B)** Western blot detection of LC3B-II (16 kDa) upon treatment with Gal-9 (300nM) or chloroquine (CQ, 50μM) for 6h. **(C)** Quantification of the fold increase in **(C)** LC3B-II induced by Gal-9 and **(D)** basal autophagic flux induced by CQ in a panel of AML cell lines with different sensitivity to Gal-9. **(E)** Flow cytometry-based cell counts of AML cell lines treated with a dose range of Gal-9. **(F)** as in **(E)** for CQ. **(G)** Flow cytometry-based cell counts of patient-derived CD34^+^ AML cells treated with Gal-9 or CQ. **(H)** Cell viability as determined by the MTS assay of U-937 cells treated with Gal-9 (300nM) in combination with a dose range of AraC upon 72h of incubation. Patient-derived AML samples were classified as **(I)** AraC responders or **(J)** AraC non-responders based on their sensitivity toward AraC as determined by flow cytometry-based cell viability upon 72h of incubation, being a non-responder when staying above 80% of cell viability even at the highest tested dose AraC. Cell viability (using MTS assay) of the parental vs. AraC resistant AML cell line panel using a low dose Aza (2,5μM) and a low dose AraC, being **(K)** 200nM for U-937, **(L)** 750nM for HL-60, **(M)** 20.000nM for THP-1 and **(N)** 750nM for MOLM-13. Of note, Aza was pre-incubated for 16h before adding Gal-9 and incubated for an additional 72h.

