## Supplementary material for "Galectin-9 has non-apoptotic cytotoxic activity towards Acute Myeloid Leukemia independent of cytarabine resistance": Suppl Figure 1

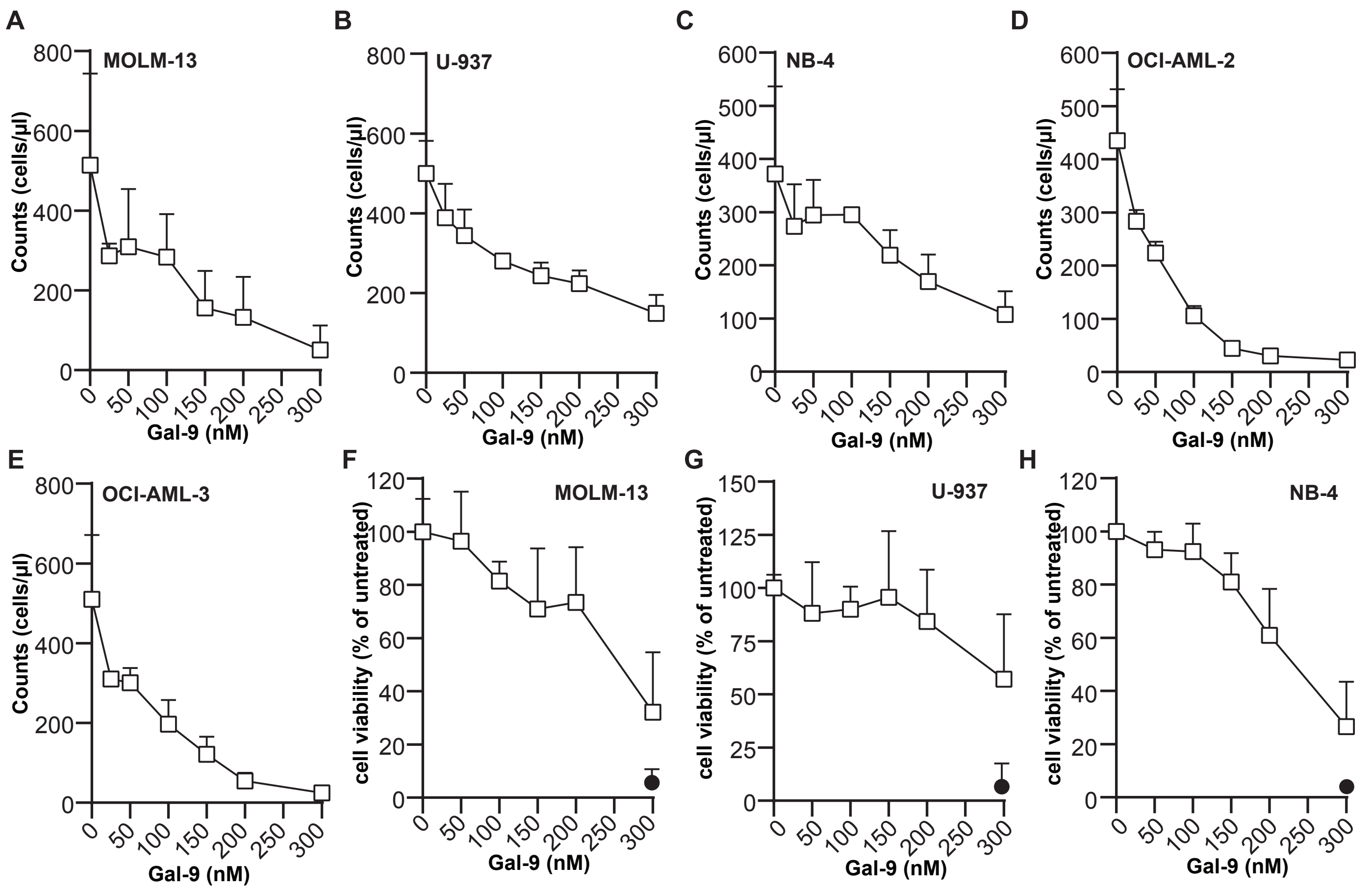


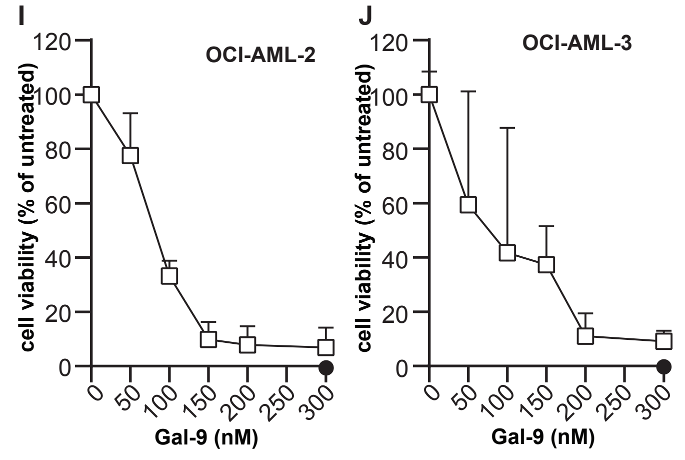


K

| EC50 | Counts | MTS | |
|---|---|---|---|
| U-937 | 193 ± 37 | >300 | High EC50 |
| NB4 | 211 ± 36 | 190 ± 30 | High EC50 |
| MOLM-13 | 114 ± 12 | 244 ± 17 | High EC50 |
| HL-60 | 135 ±32 | 172 ± 32 | High EC50 |
| OCI-AML-3 | 105 ± 43 | 142 ± 15 | Low EC50 |
| THP-1 | 104 ± 2 | 115 ± 12 | Low EC50 |
| OCI-AML-2 | 92 ± 15 | 120 ± 12 | Low EC50 |

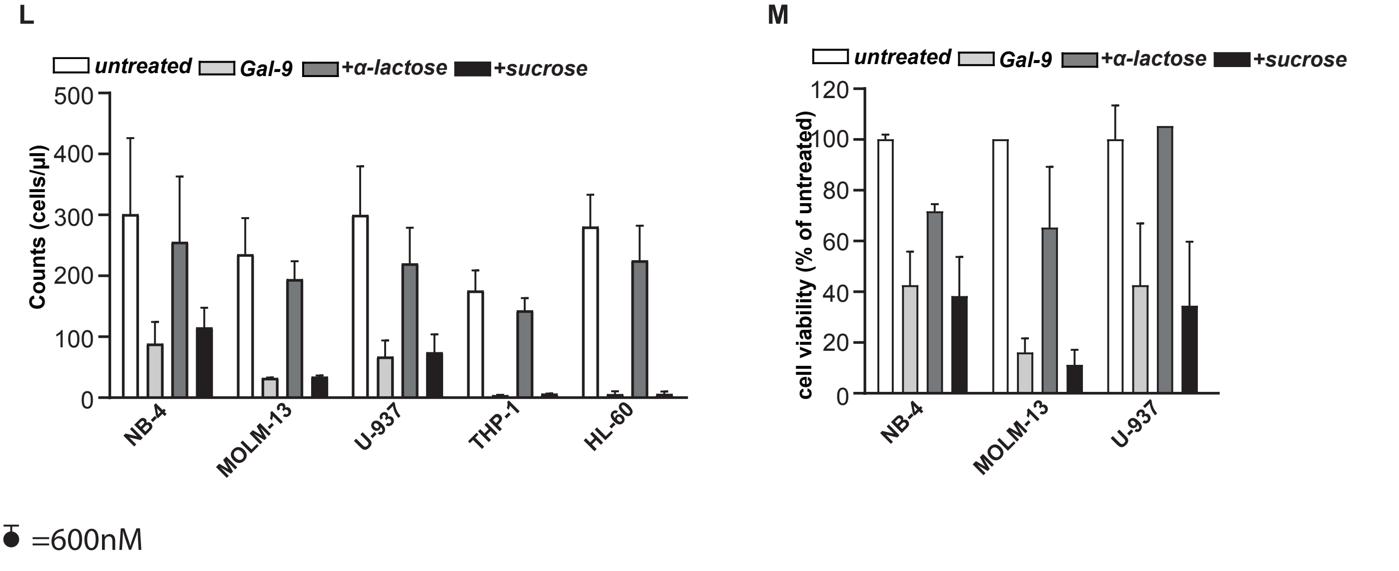


● =600nM
