## Supplementary figures and images for "Galectin-9 has non-apoptotic cytotoxic activity towards Acute Myeloid Leukemia independent of cytarabine resistance"

### Suppl Figure 2

#### Suppl Figure 2

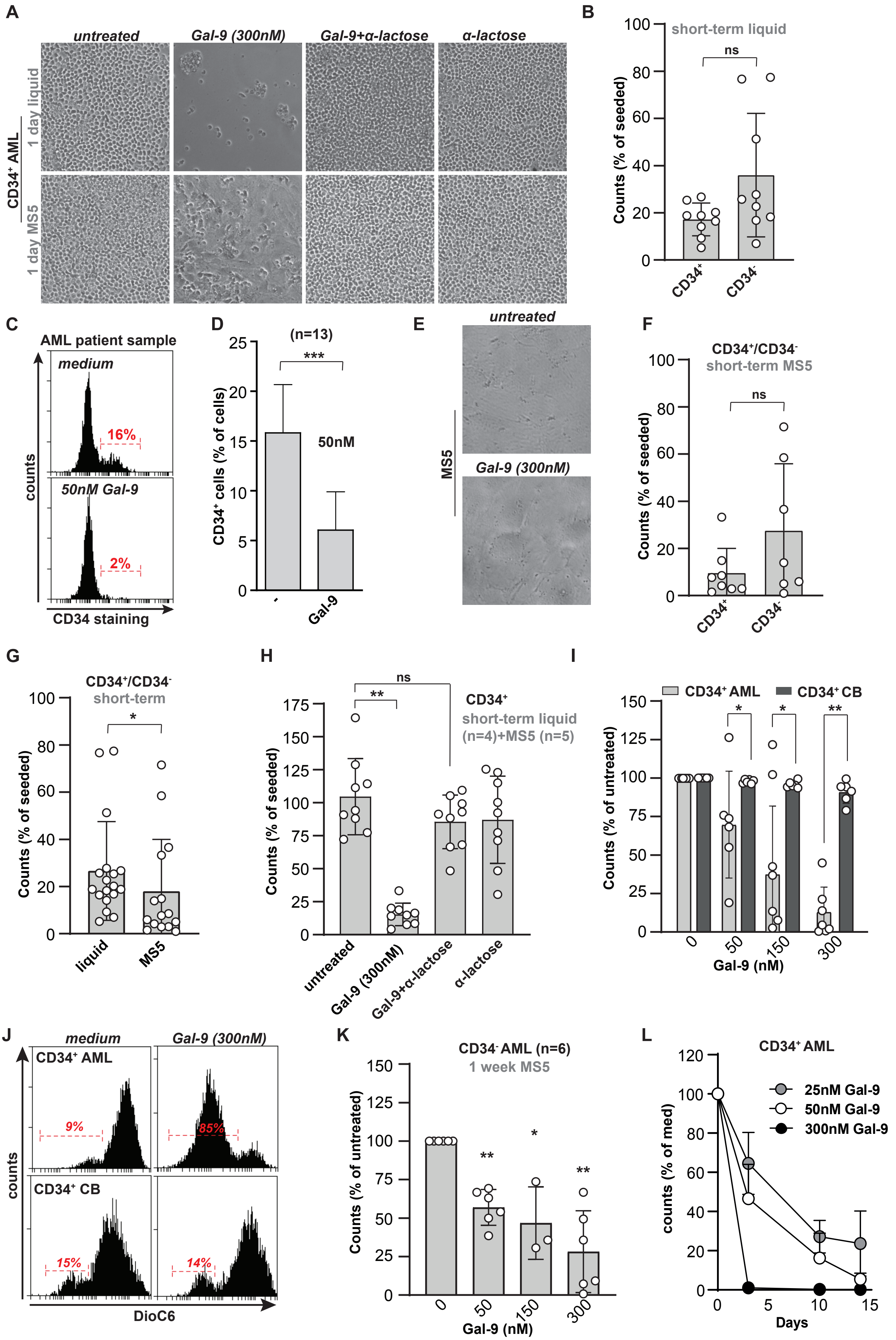

### Suppl Figure 3

#### Suppl Figure 3

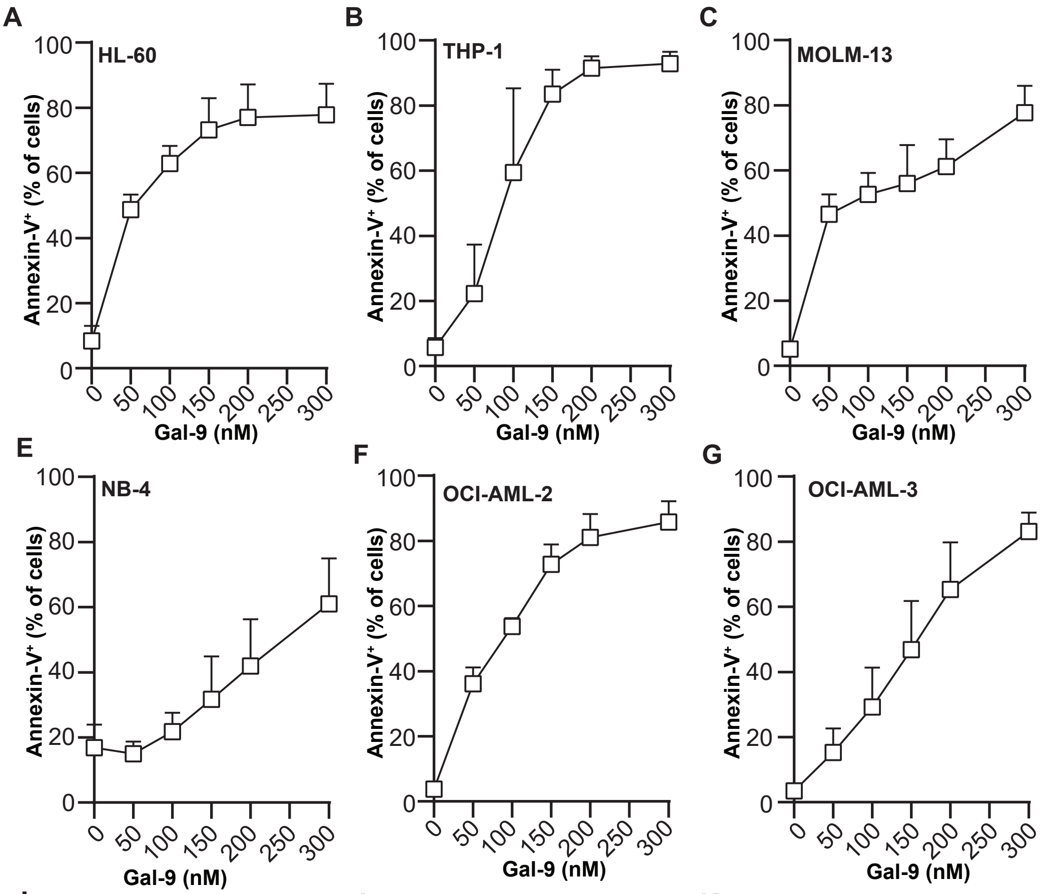


H

| *EC50* | AnnV |
|---|---|
| HL-60 | 77 ± 17 |
| MOLM-13 | 86 ± 39 |
| U-937 | 140 ± 21 |
| THP-1 | 103 ± 11 |
| NB4 | 128 ± 31 |
| OCI-AML-2 | 79 ± 20 |
| OCI-AML-3 | 126 ± 41 |

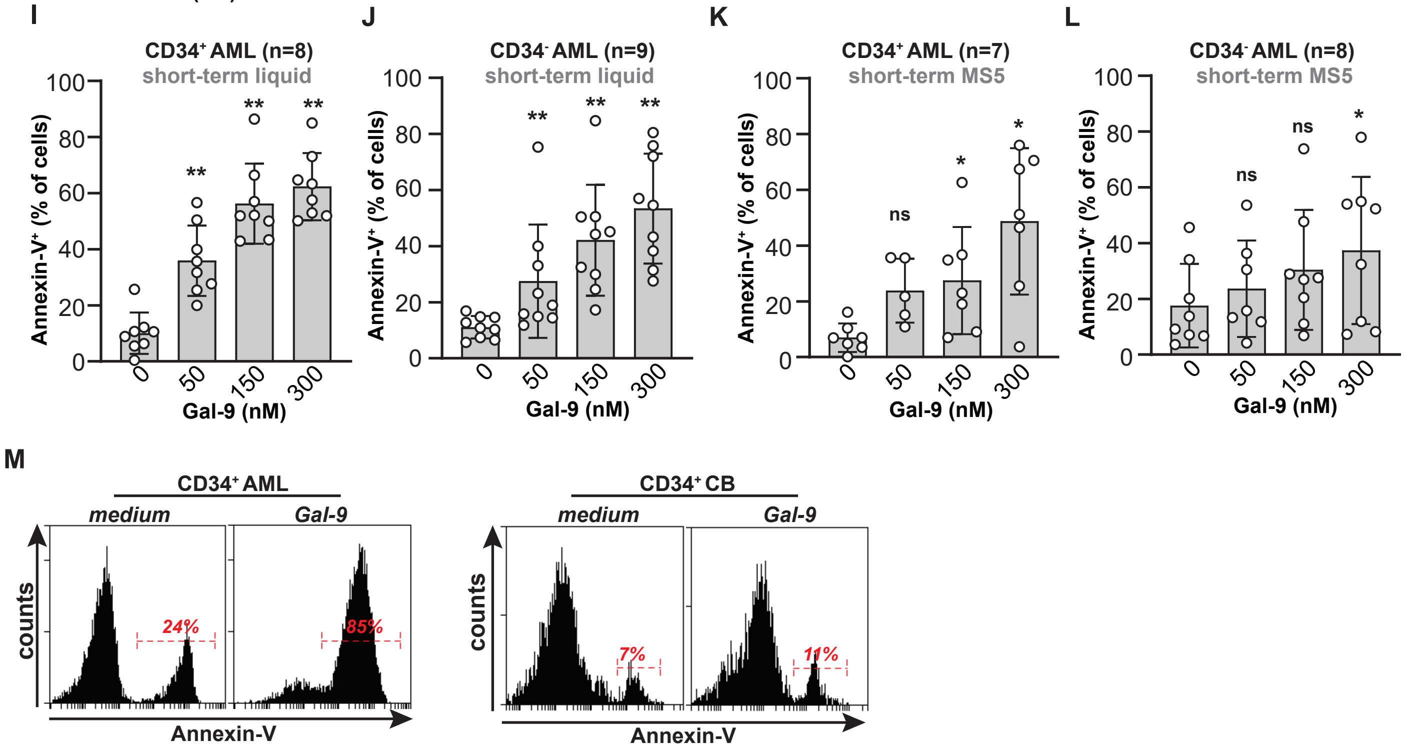

### Suppl Figure 4

Suppl Figure 4

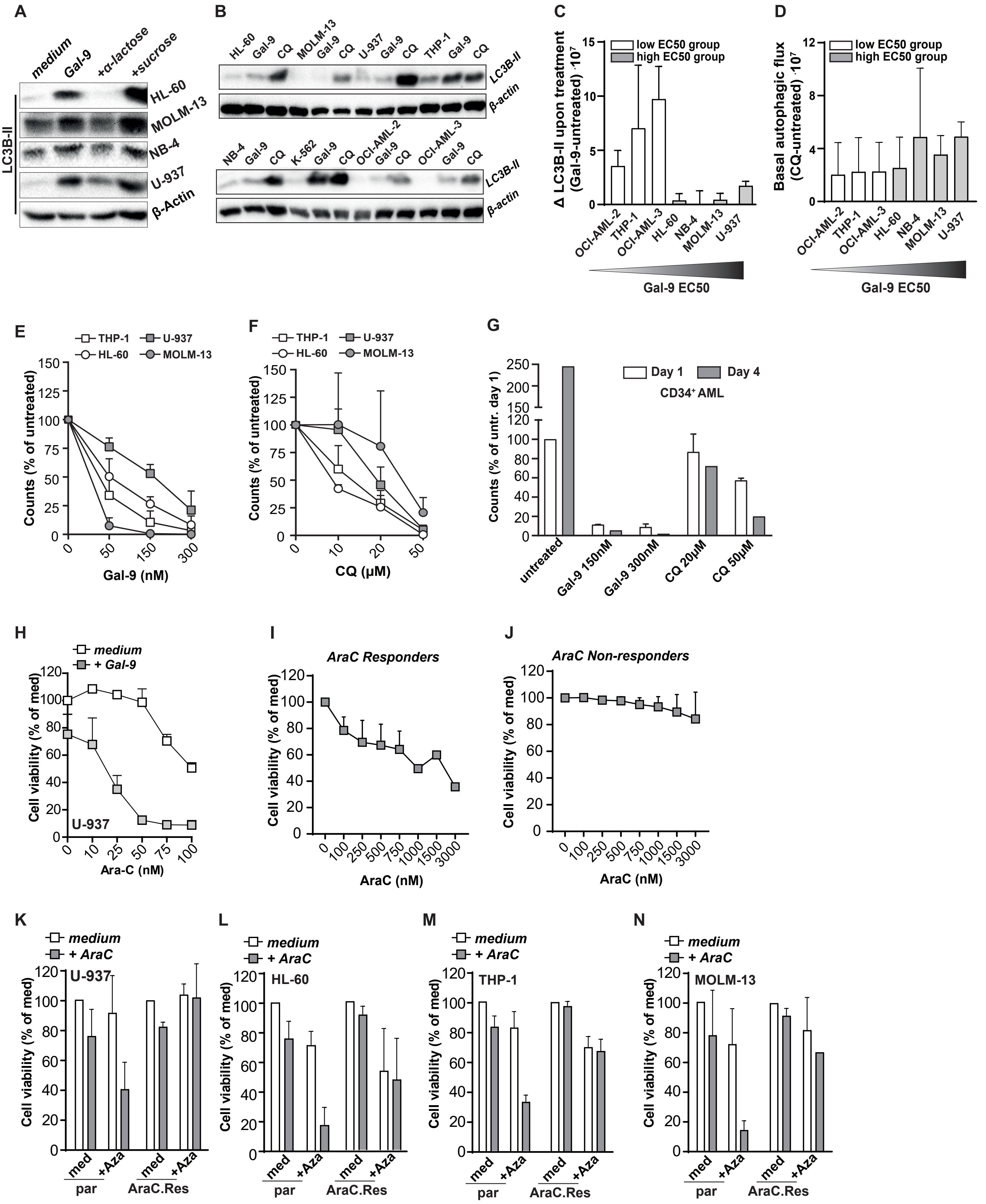
